# Facilitators and Barriers to Linkage to HIV Care and Treatment among Female Sex Workers in a Community-based HIV Prevention Intervention in Tanzania: a qualitative study

**DOI:** 10.1101/674077

**Authors:** Daniel Nyato, Soori Nnko, Albert Komba, Evodius Kuringe, Marya Plotkin, Gaspar Mbita, Amani Shao, John Changalucha, Mwita Wambura

## Abstract

**Background:** HIV-infected female sex workers (FSWs) have poor linkage to HIV care in sub-Sahara Africa.

**Methods:** We conducted 21 focus group discussions (FGDs) to explore factors influencing linkage to HIV care among FSWs tested for HIV through a comprehensive community-based HIV prevention project in Tanzania.

**Results:** Influences on linkage to care were present at the system, societal and individual levels. System-level factors included unfriendly service delivery environment, including lengthy pre-enrolment sessions, concerns about confidentiality, stigmatising attitudes of health providers. Societal-level factors included myths and misconceptions about ART and stigma. On the individual level, most notable was fear of not being able to continue to have a livelihood if one’s status were to be known. Facilitators were noted, including the availability of transport to services, friendly health care providers and peer-support referral and networks.

**Conclusion:** Findings of this study underscore the importance of peer-supported linkages to HIV care and the need for respectful, high-quality care.

## Introduction

Linkage to care, defined by the World Health Organisation (WHO) as the first HIV-specific clinical visit (1), is critical for initiation of HIV-related medical, psychological and social services for individuals newly diagnosed with HIV infection. Delays in linkage are associated with lower levels of viral suppression, greater likelihood of viral resistance, increased HIV morbidity, mortality and transmission (2–4). In Tanzania, the HIV Impact Survey (THIS) found that only about 52% of HIV positive adults age 15 and older in the general population had viral load suppression, in part because of delayed or low linkage to care (5).

HIV is not uniformly distributed across populations. Given the individual, network-level and structural risk determinants of HIV, key populations such as female sex workers (FSWs) face a disproportionately high risk of HIV infection (6). In Tanzania, FSWs are among the priority populations for expansion of HIV prevention and care services (7). Despite a shift in HIV testing strategy (8), access to HIV treatment and care services among FSWs remains limited (9). Studies conducted elsewhere in SSA have highlighted barriers to linkage to care and treatment among female sex workers including discrimination by hospital staff (10, 11), lack of money for transport (11), and lack of knowledge about HIV treatment centres (12).

Sauti program is a PEPFAR-funded program run jointly with the Ministry of Health, Community Development, Gender, the Elderly and Children (MOHCDGEC) through USAID. This community-based HIV combination prevention project, which has been operating in Tanzania since 2015, offers clinical and structural support services to key and vulnerable populations in 14 regions. Services are offered to Sauti beneficiaries through mobile outreach units, operating in hot spots such as truck stops, brothels, bars and mining centres. To ensure improved linkages, Sauti project offered transport support or escorted referral to all FSWs newly diagnosed with HIV infection to nearby health facilities. Peer educators (PEs) and home-based care providers (HBCs) employed by civil society organisations (CSOs) help FSWs’ linkage to care through escort to the health facilities. Despite these strategies, not every FSW diagnosed with HIV infection was successfully linked to HIV care.

We employ a social-ecological perspective to explore more precisely what factors prevented or facilitated FSWs testing positive in the Sauti Project services to initiate HIV care, with the aim of adapting programs to increase initiation rates. Indeed, a study conducted in Mbeya found that about a third (31%) of individuals testing HIV positive did not link to care for a period of six months (13). An exploration of the barriers and facilitators of linkage to care among FSWs is critical for designing effective interventions to reach national and global goals for HIV control.

## Materials and methods

This exploratory qualitative study was conducted in February and March 2017 – the second year of Sauti Project implementation in Tanzania. We conducted participatory group discussions in four regions (Dar es Salaam, Mbeya, Iringa and Shinyanga). These regions were selected because they were among the first regions where Sauti Project rolled-out, had a high number of FSWs compared to other regions in Tanzania (14), and had relatively high HIV prevalence (15). In the study regions, fieldwork took place in wards that were considered of high HIV risk especially those with mines, plantations or highways.

### Sampling and data collection

Snowball sampling was used to recruit participants into a group discussion. This strategy was employed due to the highly stigmatised and criminalised nature of sex work in Tanzania. In each region, CSOs staff introduced researchers to three FSWs receiving HIV testing though Sauti Project. These women received information about study aims and procedures and were asked to invite up to three other FSWs receiving Sauti services. Upon reaching a group of 10 – 12 participants, the group discussion was conducted. Criteria for participation included being beneficiary of Sauti services, living in the study area, being 18 years or older, reported having received money for sex at least once within the past three months, and provided consent to participate. The main defining socio-demographic characteristics collected were age, education and the venue where individuals’ solicited clients. Following data saturation, 227 FSWs participated in 21 focus groups.

### Procedure

All participatory group discussions were conducted in venues provided by CSOs. The discussion sessions lasted for about 90 – 120 minutes, were moderated by trained researchers and were audio-recorded. Participants were familiar with these settings as CSOs provided HIV-related services including HIV testing and family planning services. Before the beginning of fieldwork, graduate research assistants (RAs) were trained on the protocol, discussion guide and principles of research involving human subjects and informed consent. All RAs were Swahili native speakers and had experience conducting qualitative studies among key and vulnerable populations including FSWs. RAs built rapport with study participants and assumed neutrality by distancing themselves from program implementers (i.e., they declared to the participants that they were researchers and not part of the Sauti Project implementers).

A qualitative participatory group discussion guide with open-ended questions was used to facilitate discussion sessions. The discussion guide covered personal experiences with HIV testing services before and during access to Sauti Project services. Themes in the guide included factors that influenced the use of HIV testing and reasons for accepting or avoiding linkage to HIV care facilities.

We conducted one participatory group discussion in Dar es Salaam to determine the suitability of the questions asked and practical issues with conducting discussion sessions with FSWs. After it was clear that no significant changes were needed in the guide or procedure, thus the data collected was included in the analysis. During data collection, debriefing sessions were held between research assistants and senior researchers after every group discussion session to discuss the depth of the data collected and emerging issues. We obtained written informed consent from all participants prior to beginning the group sessions. To protect the identity of the participants, we asked them to use pseudonyms during consenting. Numbers were assigned to every participant for reference during discussion.

### Data analysis

The audio-recorded interviews were transcribed verbatim. All transcripts were translated into English, entered into QSR NVIVO 10 software (16)and coded by two researchers involved in data collection. A pragmatic approach to analysis was adopted, whereby coders used predefined (anticipated) codes and grounded codes. The predefined codes were developed from the research objectives, prior knowledge and repeated reading of data in early stages of analysis. These codes were further refined through further reading of the data. Grounded codes were developed through a thorough reading of the data by the two researchers in consultation with the data collection team. These codes reflected participants’ own language in ways they expressed their ideas. Based on these codes, more conceptual categories were developed and finally into themes. Widely shared views supporting emerging theories were examined alongside deviant cases. In the results, we also present deviant cases as appropriate. Representative quotes illustrating the main findings were identified from various groups.

### Ethical considerations

The Medical Research Coordinating Committee (MRCC) of the National Institute for Medical Research (NIMR) in Tanzania (NIMR/HQ/R.8c/Vol.I/432), and Institutional Review Board of the Johns Hopkins University (IRB00006985) approved this study. Approval to work in the study communities was obtained from respective local government offices after authorisation from the regional and district government authorities. All participants completed and provided written informed consent prior to beginning the group sessions. Since sex work is illegal in Tanzania, participants were asked to use pseudonyms during consenting to protect their identity. Numbers were assigned to every participant for reference during discussion. Researchers clearly explained that participation was voluntary and that the participant can decline to participate at any time if she wished to do so. Every attempt was made to minimize biases and ensure the study was conducted in the most ethical manner possible.

## Results

### Socio-demographic characteristics

Women who participated in the study were above 18 years old. Over half of the study participants (58.6%) were aged 18 – 23. In terms of education, 59.5% had attained at most primary school education, and the majority reported to solicit their sexual partners through venues, especially in bars, brothels and homes (67.4%). The distribution of ages, education levels and sites where FSWs sought clients are presented in table 1 below.

**Table 1:** Characteristics of study participants.

| Factor | Values/Categories | N = 227 (%) |
| --- | --- | --- |
| Age | 18-23 years | 133 (58.6) |
|  | 24+ years | 94 (41.4) |
| **Education** | Primary school or less | 135 (59.5) |
|  | Secondary school or above | 80 (40.5) |
| **Sites** | Street | 48 (21.1) |
|  | Facility-based (bars, brothel & home) | 153 (67.4) |
|  | Phone/internet | 26 (11.5) |

### Facilitators of linkage to HIV care among FSW

Participants discussed several factors influencing FSWs’ decision to link to care in the Sauti Project implementation sites.

### The role of peer-educators

Participants identified peer educators as the primary source of information on available HIV care services through the Sauti Project and facility-based HIV services. According to participants, availability of peer educators encouraged FSWs to go for HIV testing, and it influenced their decision to accept a referral to an HIV care clinic for further services. FSWs reported that when someone is sick, she usually seeks advice from peers about the illness and the peers recommend where to go for medical consultations. For example, the types of services offered, where to get them, whom to contact for help at times of need, and benefits of early initiation of ART.

Peer-escorted referral to treatment services was an essential influence on FSWs’ decision to accept their first HIV clinical visit to the health facilities. Participants indicated that escorted referral helped clients to receive the service quickly, and made them feel cared. FSWs pointed out that escorted referral increased their confidence and helped to reduce fears, especially during the first HIV clinic visits. For example, in one PGD a participant described that:

> *The good thing is that the home-based counsellors and peer educators knew the service providers, so before you go to the facility, they have already talked to the providers and you get the service immediately. However, when you go alone*, *you get many challenges; first, you ask yourself how should I ask about this HIV service … [laughs] … there are many nurses and doctors there…you may end-up leaving without seeing them [laughs] … if you go alone you wait for a long time because no one knows you and you are afraid to ask* – **PGD_2**

### Facilitation of transport during the first HIV clinical visit

Participants applauded about transport facilitation to attend the first HIV clinical visit for FSW to minimize some inconveniences e.g. financial constraints to seek services. Participants reported that sometimes FSWs who may have tested HIV-positive or have doubt about their HIV status, fail to present themselves to an HIV care clinic for further HIV services because they cannot afford the transport costs. To them, the facilitation of transport to the clinic was an important motivation for linkage to care.

### Presence of project-linked FSW-friendly health care providers at the health facilities

Participants pointed out that the friendly provider-client relationship had a significant impact on linkage to care. Most participants reported that the presence of service providers trained by Sauti Project to provided FSW-friendly services and motivated FSWs to attend HIV care services. According to some FSWs, health workers who received training from the programme were ‘less stigmatising’, ‘knowledgeable about FSWs’ health needs’, ‘approachable’, and ‘confidential’. The importance of the presence of friendly health care providers was emphasised in all PGDs. In one PGD a participant said:

> *For us (FSWs), it does not matter where you get the service (HIV care services) … what matters is … does the nurse or doctor understand me? How does she/he look at me? If I tell my stories about my clients and my situation [infection], is she ready to treat me nicely and will not tell other people that I have the disease [HIV]? –* **PGD_18**

### Availability of peer support network

Community engagement through peer educators was a powerful motivator for many FSWs to seek HIV related information and services. Some participants recounted a story about the feelings of social isolation and loneliness when someone receives the HIV-positive diagnosis. “*Testing HIV-positive makes you feel as if you are left alone, and that everyone leaves you because of your condition*” **[PGD_4].** Participants pointed out that, the presence of the Sauti Project and interaction between HIV positive FSWs with peers who had similar situation minimized fear of isolation and revived hopes of living like others even after an HIV-positive diagnosis. Usually, HIV positive FSWs felt less alone and less stigmatised when they met other FSWs with the same HIV status. HIV positive FSWs felt to be part or integrated into the networks of people living with HIV, thus willing to be escorted by their peers to the health facility for treatment and care. A participant during the discussion pointed out that:

> *“The advantage is that the one who tells you go to the health facility has been through the same situation and the same feelings you have, so it becomes easier to accept because you know that whenever you have a problem, she can guide you on what to do. Also, they know each other; encourage one another … so you are not alone any more”* **- PGD_1**

### Barriers to linkage to HIV care and treatment

The analysis identified barriers to linkage to HIV care that applied to FSWs in the Sauti Project implementation sites. Overall, barriers to linkage to HIV care are presented and discussed in multiple levels namely; system-level, societal-level and individual-level factors framed participants’ discussion on barriers to linkage to HIV care.

### System-level barriers

#### Fear of breach of confidentiality

All participants perceived referral to be the services offered to individuals diagnosed with HIV either through peer escort or given self-referral to the nearby health facility for HIV care registration and initiation into HIV care and treatment. Subsequently, in all PGDs, participants reported fear of breach of confidentiality as one of the significant barriers for FSWs diagnosed with HIV to enrol to the health clinic for care and treatment services. FSWs indicated that some healthcare providers working at the clinics “tell other people about their clients’ HIV status”. It was also reported that some of the health workers who were aware of individual positive status “alert their friends not to have sexual relations with the infected FSWs”. Participants in one of the PGD sessions echoed this concern:

> *“… male healthcare providers are leading gossipers … after attending to you (s) he will start to caution his close male friends to take care, … not to approach us, and later everybody in the village will know it [our HIV status]”* – **PGD_14**

Similarly, some participants had mistrust toward some peer educators (PEs) and home-based care (HBC) providers who were instrumental in linking clients to the HIV care clinics. In particular, participants complained that some PEs and HBC providers working with the project disclosed their status, which led to bad naming (called ‘prostitute’) and even discrimination by other community members. Discussing this barrier, some participants pointed out that:

> *[…] Truly, they [PE and HBCs] help us*, *but some after they have sent you to the health facility you will hear the news about your HIV status everywhere and […] they [community members] will start to refer you as a prostitute”* – **PGD_8**

> *“some…not all…if they escort you to the hospital, they end up spreading the information to other people that you have the virus”* – **PGD_21**

### Prolonged pre-enrolment period

Participants reported that if one accepted to be linked to HIV care, she was supposed to attend up to three sessions of adherence counselling before enrolment. Most participants felt that attending three days’ counselling sessions alerted other community members about their HIV status. Moreover, attending all those sessions needed transport cost and time that could help with something else. Some participants cited cases where clients had attended one or two sessions and decided to abscond. A remark made by a participant in one PGD session explains this sentiment:

> *“We know of people who were referred to HIV care clinic, but after realising that they will have to attend three days’ training with other HIV-positive people, they decided to abscond […]. Some do attend [the training] for one or two days […] you know the nurses don’t tell you when you will start getting the medicine, so you go there the first day, you go to class then you are told to come back next day – there is no clear information”* – **PGD_10**

#### Vertical and unintegrated HIV service delivery approach

Participants reported that some health facilities had dedicated rooms or blocks to attend for testing and counselling, enrolment to care, and administration of ART. According to participants, people who attend the facility for other general health services could easily see people living with HIV visiting those rooms. Providing services in dedicated venues were perceived to be stressful since it minimises privacy. One FSW who was against offering HIV care and treatment services at dedicated facilities away from other general services said:

> *“You normally see them [PLHA] congested at one place” … so when you go there, everybody knows that you have that dangerous disease and you become a talk of the town”*. – **PGD_5**

Many participants reported that some HIV care clinics were open for 8 hours only (from 8 am to 4 pm), and thus could not attend clients who visited the health facilities after that time. Participants suggested that the availability of health providers at the health facility all the time would help to overcome the challenges of working hours. In some districts, participants complained that some health facilities were enrolling clients to HIV care only for a few dedicated days in a week.

> *“If you go there* on *days when they do not enrol new people, they just look at you and ask you to come another day”* – **PGD_14**

### Societal-level barriers

#### Myths and rumours about ARVs

Many participants discussed widespread concerns about the side effects of ARV drugs, and that these rumours discouraged many FSWs from accepting linkage to care. Participants reported that everyone in the community knows that treatment with ARVs can cause horrifying dreams and loss of appetite. In addition, FSWs thought that ARVs could have adverse outcomes including further weakening for people suffering from opportunistic conditions. In some PGD sessions, participants expressed the belief that treatment with ART was behind the increased spate of sudden deaths occurring in the community.

> *Many people believe that if you start it (ARV) early it makes you tired and makes your body to weaken quickly. You can even die suddenly even when you are sleeping… You may feel a headache and die suddenly* – **PGD_21**

FSWs who self-identified themselves to have enrolled in treatment complained that ARVs were very powerful and required one to eat well before taking them. For many participants, the ‘strength of ARVs’ and the ‘need to eat well’ caused fear among FSWs who tested HIV positive but who had not started treatment with ART. Following this belief, participants thought it was wise for FSWs to choose whether to take the drugs despite the anticipated adverse outcomes.

> *“there are times when the family go to bed without food, and now you will know what to do with your medicine … if that is not death”* - **PGD_9**

#### The stigma associated with receiving HIV care

Almost all participants mentioned HIV stigma as a key barrier for FSWs to successful link to care. Some participants reported having experienced stigma, particularly when they interact with health workers. There were, for example, cases where FSWs reported to have been delayed services or called bad names when visiting health facilities. Participants indicated that it was also common for peers and other community members to use derogative terms and naming toward people living with HIV. According to participants, people who utilised HIV care services were referred to using stigmatising and derogatory terms such as *‘Maiti mtarajiwa’ [lit. expectant corpse], ‘Maiti anayeishi’ [lit. a living corpse], ‘moto’, [fire]* and *‘Maiti inayo tembea’ [lit. walking corpse]*. These had a negative connotation on their day-to-day life and their work as FSWs since it could cause them to lose clients.

HIV stigma did not only come from society, but also from close relatives and family members. Participants provided various examples to explain the family reactions after finding out that FSWs were using HIV medication:

> *…she [FSW] started using the medicine [ARVs] immediately after she tested HIV-positive … but because she believed they [family] would support her, she told her mother that she is using those [ARVs]. Instead of offering support, her mother called her aunt [FSW’s aunt] to tell her that the girl brought bad luck in the family and that she was a prostitute, and everyone who came home was told the same things […] even if it were you, would you accept to start the medicine?* – **PGD_16**

With multiple sources of stigma, participants discussed that some FSWs do not accept to go to HIV care clinics located within their villages. A few discussed that some FSWs avoided using transport offered by Sauti for clients to go to a nearby health facility to be linked to care. They instead travelled to other places to get the medicines and use them secretly.

### Individual-level barriers

#### Perceived health status

Low HIV risk perception plays a role in FSW’s acceptance of linkage to care. Those who had not developed symptoms of HIV considered themselves as being healthy, thus they did not see the importance of linking to care even when they were told that they were infected. Participants expressed that most FSWs who had taken HIV-test out of curiosity and found to have HIV infection, denied the test results because they felt that they were in ‘good physical health’. In one FGD, a young woman stated that:

> *“…You feel well just like everyone … so you continue with your life”* - **PGD_16**

Contrary to popular expressions, however, a few participants felt that these sentiments were a result of limited awareness on the mechanisms through which HIV progresses and the importance of early initiation of care. For them, the HIV virus can stay in the human body for several years while multiplying, and that manifestation of the symptoms are the final stages that you are about to die.

#### Fear of spoiling the relationship with partners or losing the clients

Some participants reported that FSWs fear to link to care as this would alert their potential clients and they would lose the clients. For those who had permanent partners feared to scare them, this would lead to separation or break the relationships. With this fear, participants reported that most FSWs found to be infected avoided to be linked to HIV care clinics because their HIV status would be known.

## Discussion

This study has documented context-barriers and facilitators for utilising HIV care and treatment. We found that some Sauti Project intervention elements were critical in influencing [motivators for] linkage to care. Consistent with other studies conducted in SSA, use of peer-educators for reaching FSWs enabled a peer-to-peer sharing of knowledge about prompt care and available alternatives for HIV services, which created a demand for both project and health facility-based HIV clinical services (17). Besides, providing support services through peer-escorted referral for individuals diagnosed with HIV promoted linkage to care. As observed in other settings, our analysis shows that peer-escorted referrals increased confidence and encouraged some FSWs to accept linkage to care (17). These findings underscore the need for expanded coverage of peer-based prevention interventions and comprehensive HIV services for FSWs (18–20). Shortcomings of a few peer educators and home-based care providers in handling clients’ information need to be addressed to increase their acceptability in facilitating linkage to care.

Whilst participants also acknowledged the role of support services such as providing transport fare and peer-support networks in promoting linkage to care, they discussed many challenges that inhibit FSWs’ linkage to HIV care at multiple levels, which merit future interventions. At a system level, breach of confidentiality and disrespect from healthcare providers; unfriendly service delivery environment; and the prolonged training sessions took before enrolment care and treatment, perceived adverse effects of ARV hindered FSWs to swiftly use care and treatment services. At a societal level, the stigma associated with accessing HIV care and myths surrounding antiretroviral treatment, prevent FSWs from accepting linkage to HIV care. At the individual level, we found that perceived health status and fear of spoiling relationships with partners, negatively impede linkage to care. Although more research is needed to address the broad range of barriers, these results highlight critical entry points for interventions to improve linkage to HIV care and treatment among FSWs diagnosed with HIV (21).

Participants described how fear of stigma and discrimination could act as a barrier to linkage to HIV care or lead to delays in deciding to visit an HIV care and treatment facility after an HIV diagnosis. FSWs often face multiple levels of stigma related to the social and structural context of sex work (19, 22, 23) or as a result of household illness (24). Thus, it is not surprising that stigma and discrimination continue and perhaps increasing among FSWs living with HIV. In our study, FSWs highlighted that stigma and discrimination in the form of verbal harassment and disclosure of HIV status within the healthcare setting were significant barriers to their linkage to HIV care, which is likely because of their sex work practices and HIV status combined. Interventions focused on health service providers to reduce stigma and discrimination, such as sex work sensitisation training and participatory dialogue, are urgently needed as they have the potential to increase linkage to HIV care among FSWs, a finding that has been highlighted in other studies (20, 25). There is also a need for community sensitisation that highlight the importance of support systems for people living with HIV, to enhance acceptance of those who are diagnosed with HIV (26). It is known that healthcare providers’ stigma towards people living with HIV can be a reflection of the broader negative social norms embedded in the community (27). Thus, addressing the drivers of HIV stigma especially attitudes towards people living with HIV and a lack of awareness on what constitutes stigma is critical to increasing linkages to HIV care. Studies have shown that interventions which empower the community are useful and effective means to increase tolerance and reducing HIV stigma (28). Therefore, it is crucial that community-based HIV interventions such as Sauti include interventions aimed at increasing tolerance and lessen HIV stigma at all levels.

Unfriendly service delivery ‘environment’ emerged as a leading barrier for FSWs to initiate visits to HIV clinics. Although studies have shown that having a dedicated section for HIV services at the facility level may increase the quality of care (29), this study found that FSWs were concerned that such arrangement exposes them to other patients attending the facilities, thus increasing individual stigma. In addition, while studies show that the length of adherence sessions increases adherence to treatment in the general population (30), FSWs in this study raise this as a concern and that it discourages them as it compromises their working time, and that some health facilities opened only for eight hours a day. These findings point to the need for communicating thorough information to FSWs on what to expect when they visit the clinic, including whom they will see and when, and if possible the average time they should expect to spend at the clinic for each visit and their rationale (31). Besides, scepticism regarding the location of the clinic-space and the hours of operation shows the pervasive nature of anticipated stigma, especially the expectation that when other people see them will gossip, point fingers, or discriminate them (27). The findings further point to the potential for initiation of pre-ART outreach services through FSWs’ existing networks. It is possible to use peer-educators to organise FSWs’ in teams within their villages for pre-ART counselling sessions and sample collection for confirmatory tests and other clinical procedures. Community-based ART evaluations have shown that initiation of care through outreach service has the most significant effect on the timely initiation of care, which leads to positive health outcomes (32).

The myths and rumours about ARVs, coupled with a lack of correct information on HIV and disease progression were barriers to linkage to HIV care. In this study, participants discussed that the use of ARVs caused deterioration of the body and was the leading cause of the observed sudden deaths in the community. These myths created fear among those who are newly diagnosed with HIV infection and an increasing tension to those who are on treatment.

The continued myths and rumours about ARV points to the need of robust community programme and a comprehensive communication plan, which will address how HIV stigma plays out through multiple modalities as suggested in other studies (33). Such a programme must ensure that there is a regular, open and consistent dialogue within the community, and at all local government structures for a sustained information sharing. Local tailoring of messages and materials may be the best approach for addressing the myths and rumours preventing FSWs from linking to HIV care.

It is also important to provide more information regarding the health benefits of timely initiation of ART to individuals diagnosed with HIV, and the preventive effect on the spread of HIV to sexual partners when viral suppression has been achieved (34). Findings from this study show that some FSWs fear linkage to care because their partners will know that they are HIV-positive and are using ART, which can spoil their relationship. This suggests a lack of understanding of the preventive benefits of early initiation of HIV treatment and the need for partner involvement in HIV testing and counselling through index testing. Right messaging to show that individuals who are virally suppressed not only are less likely to develop HIV-related complications but also are less likely to pass on the virus to sexual partners (35). Availing this knowledge to the community can reduce the enacted and perceived stigma, which inhibits many people from disclosing their HIV status and seeking appropriate care.

HIV risk perception, e.g. lack of HIV signs and symptoms consistently emerged as a barrier to linkage to care. This finding suggests limits in community awareness of HIV progression and the importance of early initiation of ART. Such knowledge is urgently needed if programs are to increase FSWs accepting to be initiated to HIV care. Results in this study also found that FSWs are more likely to trust their peers and rely on their informal networks to provide them with information on available health services. Since FSWs are hard to reach, it could be a challenge for healthcare providers to reach and deliver the needed health information necessary to change the risk perceptions hindering linkage to care. However, studies have shown that in contexts where sex work is illegal, most FSWs engage in some business that helps to guise their identities. Such business includes being a bar attendant, working in massage pallor, food selling spaces, saloon and so on (36, 37). Reaching workers in these places with health information, and recruiting peers who would reach FSWs will increase the reach of health literacy on the importance of prompt HIV care, which will have the potential to increase linkage to care. The results from previous studies on the effectiveness of peer education programs for FSWs showed that peer education intervention significantly increased knowledge on HIV (38), reduced STIs, and increased condom use among FSWs (39). Thus, interventionists and health professionals may achieve the goal of increasing linkage to care by working closely with peer educators.

Other studies conducted in SSA have also underscored how transport related costs could hinder linkage to care (40–42). Consistent with those studies, our study underscores the need to pay attention to salient issues such as logistical constraints that can negatively influence FSWs decision on linkage to HIV care.

### Study limitations

As study participants were limited to FSWs who received services provided by Sauti Project, the information may not describe barriers and facilitators to linkage to HIV care for FSWs in non-Sauti implementation areas. Readers ought to be cautious that participants in this study were not selected based on HIV status, it is likely that some of the information shared is based on perceptions and not lived experiences. Future studies may want to interview FSW stratified into groups based on HIV status if this is feasible and can be done in a non-harmful way.

## CONCLUSION

Linking HIV-positive FSWs to care is complex and involves a range of barriers at the system, societal and individual levels. Although the socio-ecological perspective provides an approach for understanding how these multi-level factors interact to influence linkage to care, this study underscores the need for understanding the salient factors for program prioritisation. In addition, health services and future interventions should consider the salient facilitators and barriers identified in this study to improve linkage to care among FSWs in similar contexts.

## Acknowledgements

We thank the women who volunteered to participate in this study. A debt of gratitude is also owed to research assistants, civil society organizations and peer educators working in study regions. We appreciate leaders of FSW networks, CSOs working with FSWs and local government authorities, especially the village leaders and ward executive officers in areas where this study was conducted for their support and guidance.

